# Protein interactions of *Magnaporthe oryzae* protein kinase CK2 and secondary data analysis of a large number of transcriptomes suggest chaperone-like activity is integral to its function

**DOI:** 10.1101/704775

**Authors:** Lianhu Zhang, Dongmei Zhang, Dan Liu, Yuan Li, Hongchen Li, Zonghua Wang, Bjoern Oest Hansen, Stefan Olsson

## Abstract

CK2, a serine/threonine (S/T) kinase present in eukaryotic cells is known to have a vast number of substrates. We have recently shown that it localizes to nuclei and at pores between hyphal compartments in *M. oryzae*. We performed a pulldown-proteomics of *M. oryzae* CK2 catalytic subunit MoCKa to detect interacting proteins. The MoCKa pulldown was enriched for septa and nucleoli proteins and intrinsically disordered proteins (IDPs) containing a CK2 phosphorylation motif proposed to destabilize and unfold alpha helixes. This points to a function for CK2 phosphorylation and corresponding phosphatase dephosphorylation in the formation of functional protein-protein aggregates and protein-RNA/DNA binding. To test this as widely as possible we used secondary data downloaded from databases from a large range of *M. oryzae* experiments and also for a relatively closely related plant pathogenic fungus, *Fusarium graminearum*. We found that CKa expression was strongly positively correlated with S/T phosphatases as well as with disaggregase (HSP104, YDJ1, SSA1) and an autophagy indicating protein (ATG8). The latter points to increased protein aggregate formation at high levels of CKa expression. Our results suggest a general role for CK2 in aggregation and disaggregation of IDPs and their binding to proteins, DNA and RNA interactions.

## INTRODUCTION

CK2 is a serine/threonine (S/T) kinase constitutively active in eukaryotes (Meggio and Pinna, 2003). The CK2 holoenzyme is a heterotetrameric structure consisting of two α-units and two regulatory β-subunits (Ahmad *et al*., 2008). In yeast these units can each be of two types and are named a1, a2, b1 and b2 (Padmanabha *et al*., 1990) while several other fungi, including *M. oryzae*, only contain one CKa (Zhang *et al*., 2019). CK2 have been shown to phosphorylates a large number of proteins (Meggio and Pinna, 2003; Ahmad *et al*., 2008; Götz and Montenarh, 2016) and seems involved in many cellular processes (Götz and Montenarh, 2016). Interestingly, intrinsically highly disordered SRP40 protein in yeast (homologous to Nopp140; Insect, Drosophila and Nolc1;Mammal, Human) has been shown to become phosphorylated to varying degrees by CK2 with effects on the SRP40 conformation, binding, and aggregation properties with further effects on the diverse functions of the protein (Tantos *et al*., 2013; Na *et al*., 2018). The large number of substrates of CK2, could possibly be explained if CK2 phosphorylation destabilizes/unfolds alpha helixes in IDPs (Zetina, 2001) and CK2 activity in combination with phosphatases facilitates conformation changes and binding of IDPs.

We have previously argued that filamentous fungi are better models than yeast for investigating eukaryotic cellular functions involved in cell-cell communication when we studied CK2 localization and involvement in pathogenesis (Zhang *et al*., 2019). We showed that the *M. oryzae* CK2 holoenzyme (MoCK2) accumulates in the nucleolus, that it localizes in structures near septal pores, and assembles to form a large ring structure perpendicular to the appressorium penetration pore (Zhang *et al*., 2019).

To further investigate the general hypothesis that CK2 is especially involved in phosphorylating IDPs and proteins known to be localized at septa and in the nucleolus we made a pulldown followed by proteomics using the GFP-CKa (Zhang *et al*., 2019) as our bait to find which proteins CK2 interacts with. Since CK2 is known to interact with a large number of proteins (Meggio and Pinna, 2003) we expected to find many proteins, and we did. To aid in interpreting the results we made a functional classification of the interacting proteins. The pulldown was overrepresented for proteins known to be present in nucleoli and at septa as expected and we found that CK2 interacts preferentially with intrinsically disordered proteins (IDPs) carrying a phosphorylation motif that can destabilize alpha helix folding (Zetina, 2001).

To test this destabilization/unfolding hypothesis in a general way we decided to make an analysis of secondary data already published (Smith *et al*., 2011; Zaritsky, 2018) and downloaded a large range of transcriptomic data for *M. oryzae*. The following hypotheses resulting from the general hypothesis were tested: Sub-hypothesis 1: If unfolding-refolding activity is important for IDP aggregation then a high CKa expression should be correlated with expression of some S/T phosphatases present in the pulldown that can act as a counterpart to the CK2 S/T kinase activity. Sub-hypothesis 2: Higher general aggregation activity should lead to more aggregates of misfolded proteins, especially of IDPs. Disaggregases and ATG8 are known to be needed to remove aggregates formed from misfolded proteins (Glover and Lindquist, 1998; Wong and Cuervo, 2010; Watabe and Nakaki, 2011, 2012) and the expression of these genes should be correlated with CKa expression. The correlations implied by Sub-hypothesis 2 should also apply for other fungi. To test this hypothesis further we thus downloaded a set of secondary data for another plant pathogenic fungus, *Fusarium graminearum*, and tested if similar correlations between homologous genes apply. We found that CK2 expression correlates with several CK2 phosphatase expression supporting the first sub-hypothesis. We also found that CK2 expression correlates with both disaggregase gene expression and ATG8 in both fungi tested giving support for the second sub-hypothesis.

Our work provides evidence supporting the view that one of the main roles for the CK2 holoenzyme is as a general inducer of binding, conformational changes and aggregate formation of intrinsically disordered proteins.

## RESULTS

### Identification of potential septal and nucleolar and other substrates for MoCK2 by GFP-CKa pulldown and investigation if these proteins contain an CK2 phosphorylation unfolding motif

The CK2 localization pattern suggested that CK2 may have substrates associated with septa and nucleolar function (Zhang *et al*., 2019). To explore this, we performed co-immunoprecipitation to identify proteins interacting with CK2 using GFP-CKa as a “bait”, and in addition to the bait, identified 1505 proteins (**Data S1**). There is the risk of false positives in a pulldown. We estimated the number of potentially false positives and removed 155 (~10%) of the lower abundance proteins to arrive at a list of 1350 CKa interacting proteins (see Methods). Both CKb1 and CKb2 were found in the pulldown with GFP-CKa (Data S1) although the amounts found in the pulldown were not supporting that 2CKa are always interacting with 1 CKb1 and 1 CKb2 to form the CK2 holoenzyme (Zhang *et al*., 2019). The CK2 holoenzyme can be formed with only type of CKb subunit (Ahmad *et al*., 2008) although our previous results showed that both MoCKb subunits are needed for normal growth and pathogenicity of *M. oryzae* (Zhang *et al*., 2019).

Since CK2 localizes to septa we looked for known septal proteins in the pulldown. All previously identified proteins by Dagas et al. (Dagdas *et al*., 2012) that are involved in appressorium pore development, were found in the pulldown as was a protein annotated as the main Woronin body protein, Hex1 (MGG_02696). Since the Woronin body in Ascomycetes is tethered to the septal rim by Lah protein (Ng *et al*., 2009; Plamann, 2009; Han *et al*., 2014) we searched for a homologue in *M. oryzae* and found a putative MoLah (MGG_01625) with a similar structure as in *Aspergillus oryzae* (Han *et al*., 2014) that is also present in the pulldown. In addition to the Lah, 18 other intrinsically disordered septal pore associated proteins (Spa) were described for *Neurospora crassa* (Lai *et al*., 2012). We identified putative orthologs for15 of the18 Spa proteins in *M. oryzae* (See link from **Table 1**). Of these putative MoSpa proteins, six were present in the CKa pulldown, MoSpa3 (MGG_02701), MoSpa5 (MGG_13498), MoSpa7 (MGG_15285), MoSpa11 (MGG_16445), MoSpa14 (MGG_03714) and MoSpa15 (MGG_15226). MoSpa3, MoSpa5 and MoSpa15 also contain the CK2 phosphorylation alpha helix unfolding motif (**Data S1**). The presence of the a MoLah and several MoSpa homologues in the CKa pulldown is in agreement with the view that localization of the GFP-fusion protein (Zhang *et al*., 2019) gives a proper representation of CK2 localization. In addition, and further supporting this conclusion, the six septal pore associated proteins (SPA) are homologues for intrinsically disordered proteins that are expected to form temporary gels that are used to reversibly plug septal pores and regulate traffic through septa (Lai *et al*., 2012). CK2 could actively be involved in the gelling/un-gelling of the regions near septa to create a membraneless organelle controlling the flow through septa.

**Table 1.** Key resources.

| REAGENT or RESOURCE | SOURCE | IDENTIFIER |
| --- | --- | --- |
| Antibodies |  |  |
| anti-GFP beads | Chromotek,<br>Hauppauge, Ny, USA | GFP-Trap®_A |
| Deposited Data |  |  |
| Transcriptomic data sra/fastaq files for *M. oryzae* | Gene Expression<br>Omnibus | https://www.ncbi.nlm.nih.gov/geo |
| Genome for mapping transcriptomic data | EnsemblFungi | http://fungi.ensembl.org/Magnaporthe_oryzae/Info/Index |
| Transcriptomic data *F. graminearum* | PlexDB | http://www.plexdb.org/modules/PD_general/download.php?species=Fusarium |
| Protein ID codes and annotation (FGSG_codes for *Fusarium Graminearum* Ph1 and MGG_codes for *Magnaporthe oryzae* 70-15) | BROAD | ftp.broadinstitute.org/distribution/annotation/fungi/ |
| List of proteins with unfolding motif according to (Zetina, 2001) found for *M. oryzae* found using the FIMO tool at the MEME-suite website. | Figshare, this paper | https://figshare.com/s/a25ef9d1b93240e72f96<br>DOI:<br>10.6084/m9.figshare.7066379 |
| Nucleolar proteins in the MoCKa pulldown and annotation of *M. oryzae* with the help of *S. cerevisiae* annotation. | Figshare, this paper | https://figshare.com/s/752257b8b25dfe19a4e0<br>DOI:<br>10.6084/m9.figshare.7067897 |
| *M. oryzae* transcriptomic data matrix covering a range of experiments | Figshare, this paper | https://figshare.com/s/64eec43151576417afbc<br>DOI:<br>10.6084/m9.figshare.7068857 |
| *F. graminearum* transcriptomic data matrix covering a range of experiments | Figshare, this paper | https://figshare.com/s/e28813e5cc7b9c394e08<br>DOI:<br>10.6084/m9.figshare.7068860 |
| List of *M. oryzae* orthologs to *N. crassa* SPA proteins | Figshare, this paper | https://figshare.com/s/39c3962d1b8d7d969338<br>DOI:<br>10.6084/m9.figshare.7066382 |
| FungiFun: Funcat analysis of the *M. oryzae* CKA pulldown proteins | Figshare, this paper | https://figshare.com/s/231c764fad6e60855795<br>DOI:10.6084/m9.figshare.7066385 |
| Experimental Models: Organisms/Strains |  |  |
| *M. oryzae* Strain Ku80 transformed with the over-expressed GFP-MoCKa fusion protein (GfpA). | (Zhang *et al.*, 2019) | N/A |
| Software and Algorithms |  |  |
| SRA toolkit | NCBI | http://ncbi.github.io/sra-tools |
| FIMO tool | The MEME suite | http://meme-suite.org |
| FASTQC (RNAseq quality control) | (Andrews, 2010) | N/A |
| Functional catalogue of fungal proteins | FungiFun | https://elbe.hki-jena.de/fungifun/ |
| Other |  |  |
| Proteomics services | BGI Tech, Schenzen, Guangdong province China | N/A |

Since many other of the proteins we found in the pulldown were IDPs we also searched the *M. oryzae* proteome for proteins containing the CK2 phosphorylation helix unfolding motif identified by Zetina (Zetina, 2001) using the FIMO tool at the MEME-suit website (http://meme-suite.org/) and found 1465 proteins with the motif out of a total of 12827 proteins annotated for *M. oryzae*. We found 275 of the proteins contain at least one unfolding motif for alpha helixes (Zetina, 2001). Thus, there is an overrepresentation of the motif among the pulldown proteins (P-value for the null hypothesis of same frequency as in the whole proteome = 4 E-19, Fisher’s Exact test) lending support for the proposed role for this motif in these proteins as a target for CK2 phosphorylation and protein unfolding. The finding of overrepresentation of this signal in the set of CK2 interacting proteins corroborates the previous suggestion that CK2 is involved in the destabilization/binding of intrinsically disordered proteins (Zetina, 2001; Tantos *et al*., 2013; Na *et al*., 2018) and is consistent with the strong accumulation of both CK2 and intrinsically disordered proteins in the nucleolus (Frege and Uversky, 2015) and also at pores between cell compartments (Lai *et al*., 2012; Zhang *et al*., 2019).

To further explore the general hypothesis that CK2 could interact with and possibly phosphorylate intrinsically unfolded proteins we used FungiFun (https://elbe.hki-jena.de/fungifun/) to make a functional classification of the pulldown proteins with a special interest for those containing the alpha helix unfolding motif (Zetina, 2001). All the protein categorization classes for the pulldown is accessible at the FungiFun entry in **Table 1**. We found strong overrepresentation for proteins involved in rRNA processing among the pulldown proteins containing the alpha helix unfolding motif (Funcat category, 11.04.01 rRNA processing, p-value for same abundance as in whole genome = 1.53E-26) as well as for proteins that, themselves, are known to interact with other proteins, DNA, and RNA (16.01 protein binding *p* 1.07E-09, 16.03.01 DNA binding *p* 1.45E-06, 16.03.03 RNA binding *p* 2.77E-11, p-values for same abundance as in whole genome). These classes of proteins are enriched for intrinsically disordered proteins. Such intrinsically disordered proteins can interact with each other to form ordered subregions that have been described as membraneless organelles, such as nucleoli (Wright and Dyson, 2015). Since CK2 localizes to the nucleolus (Zhang *et al*., 2019) we were especially interested in the interaction of CK2 with nucleolar localized proteins. We identified homologues to the well described *S. cerevisiae* nucleolar proteins and found a total of 192 proteins in *M. oryzae* homologous to yeast nucleolar proteins. We found 120 (63%) of the nucleolar proteins in the pulldown and 60 of these (50% of the ones found) had the alpha helix unfolding motif (**Data S1**) (also Table 1, linked from “*M. oryzae* homologues to *S. cerevisiae* nucleolar proteins”). The nucleolar proteins were thus highly overrepresented in the pulldown (P-value for the null hypothesis of same frequency as in the whole proteome 9E-43 Fisher’s Exact test) compared to the whole proteome as was also nucleolar proteins having the unfolding motif (P-value for the null hypothesis of same frequency as in the whole proteome *p* 2E-13, Fisher’s Exact test).

Ck2 is known to interact with the disordered nucleolar protein SRP40 that has a multitude of CK2 phosphorylation sites and in addition CK2 is known to change the protein binding activity towards other proteins depending on the SRP40 phosphorylation status. When SRP40 becomes highly phosphorylated it binds to and inhibits CK2 activity in a negative feedback loop ensuring that CK2 phosphorylation level will balance (Tantos *et al*., 2013; Na *et al*., 2018). We identified a putative MoSRP40 (MGG_00613) and it is disordered in a similar way as other SRP40 proteins and it is well conserved in filamentous fungi (**Method S1**). The MoSRP40 protein was highly represented in the CKa pulldown indicating that it interacts with CKa (**Data S1**).

Interestingly, proteins that are imported into mitochondria and involved in oxidative phosphorylation (Funcat category, 2.11 “electron transport and membrane-associated energy conservation”) were enriched in the pulldown (62 in the pulldown of 130 in the whole proteome, *p*-value for the null hypothesis of same frequency as in the whole proteome 2.26E-14). In contrast to septal and nucleolar interacting proteins, the mitochondrial proteins were not enriched for the known unfolding motif.

Although CK2 has been implicated to be located in mitochondria in earlier studies in other organisms, no proteomic study of yeast mitochondria has detected the presence of CK2 (Rao *et al*., 2011). Hence, we do not expect MoCK2 to be present in mitochondria, and we saw no evidence for mitochondrial localization of GFP-CKa (Zhang *et al*., 2019). Of special interest was however the strong overrepresentation of mitochondrial proteins among the CKa pulldown proteins without the alpha helix phosphorylation unfolding motif. Since such proteins need to be imported into mitochondria in an unfolded state, this may point to the existence of CKa phosphorylation and unfolding motifs other than the one identified by Zetina (Zetina, 2001) that together with chaperone HSP70 (Yang *et al*., 2018) help keep these proteins unfolded until they reach their destination inside the mitochondria.

There was no enrichment for specific pathogenicity related proteins in the whole pulldown proteome (Funcat category: 32.05 “disease, virulence and defence”) but a significant depletion (p-value for the null hypothesis of same frequency as in the whole proteome 3.31E-17) was found. This is generally true within the main Funcat category related to stress and defence (32 “CELL RESCUE, DEFENSE AND VIRULENCE”) (p-value for the null hypothesis of same frequency as in the whole proteome 8.32E-05) with the exception of proteins involved in the unfolded protein response that was overrepresented (32.01.07 “unfolded protein response”,e.g. ER quality control) among the pulldown proteins without motif (p-value for the null hypothesis of same frequency as in the whole proteome 4.69E-04). This is notable since an involvement of CK2 in protein import into the ER has be established (Wang and Johnsson, 2005).

A strong association of pathogenicity related proteins with CK2 was not expected although deletion of any of CKb components severely affected pathogenicity (Zhang *et al*., 2019). This either because of the *in vitro* growth conditions to produce the biomass used for the pulldown or that CK2 is mainly involved in the initial stages of infection to establish the biotrophic stage (Fernandez and Orth, 2018).

To have a dynamic function as an unfolder of proteins by phosphorylation, CK2 should be partnered with phosphatases as counterparts and their activity may track CK2 activity. Interestingly, five putative S/T phosphatases (MGG_03154, MGG_10195, MGG_00149, MGG_03838, MGG_06099) were in the pulldown set of proteins (**Data S1**). Conceivably these might de-phosphorylate CKa substrates as well as substrates of other kinases to expand the reach of CK2 in regulating the phosphoproteome (Subhypothesis 1). To examine the general relationship between the expression of CK2 and these phosphatases as widely as possible, we downloaded expression data from a range of experiments with *M. oryzae* and plotted the expression of the five phosphatases found in the pulldown, and an S/T phosphatase not found in the pulldown, as a function of the CKa expression. We found that two of the S/T phosphatases (MGG_00149 and MGG_03154) present in the pulldown were strongly correlated with CKa expression and the others were less strongly correlated (**Fig. 1**) further supporting the view that CKa-dependent phosphorylation/dephosphorylation plays a major role in shaping protein interactions. Together with the high expression of CK2 in cells, this suggests an important function of CK2 as a general temporary unfolder of intrinsically disordered proteins, that comprise roughly 30% of eukaryotic proteins (Vucetic *et al*., 2003), in a similar way as it is known to interact with SRP40 and influence its activity (Tantos *et al*., 2013; Na *et al*., 2018).

**Fig. 1.**
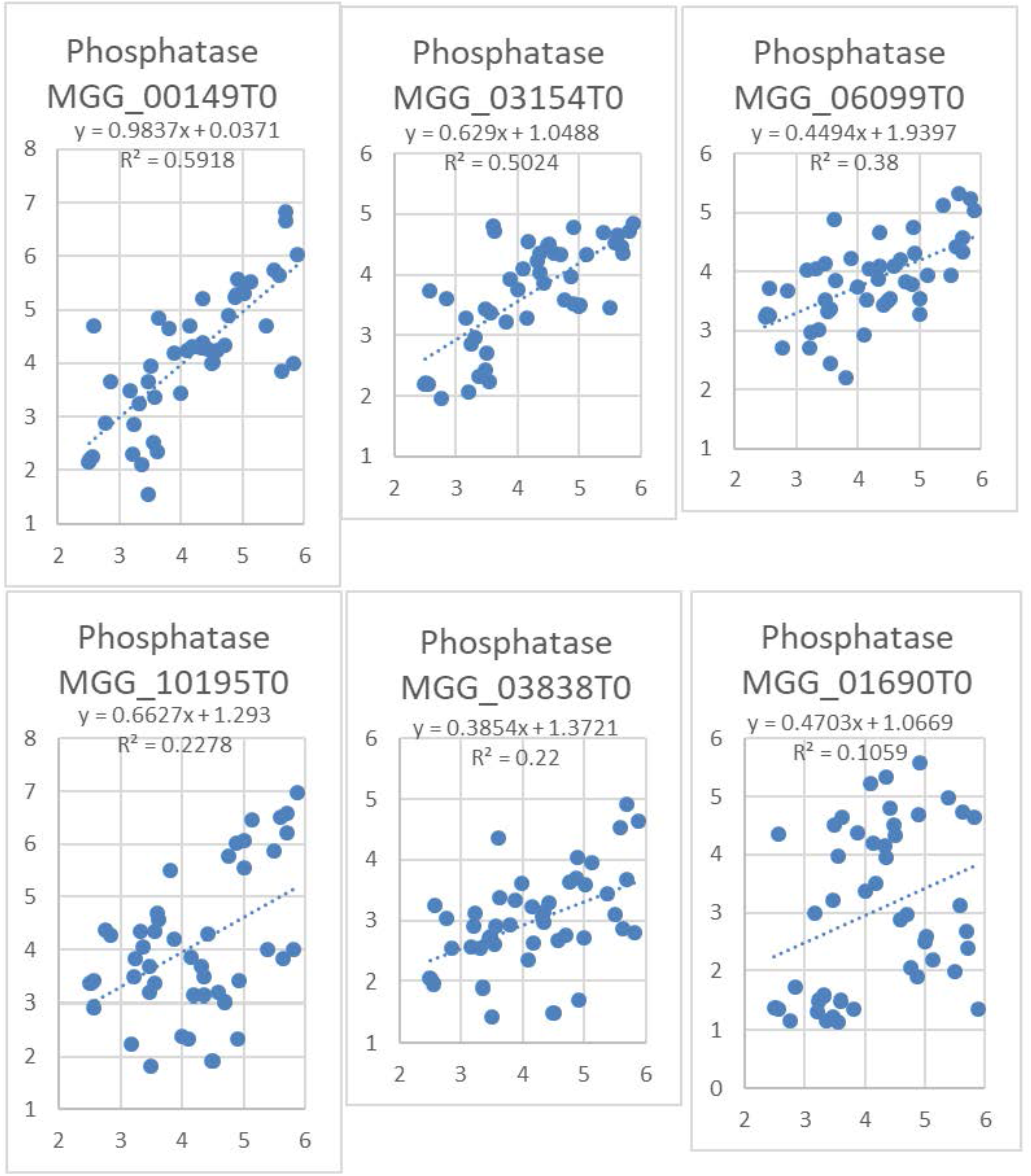
Plots of 6 putative serine/threonine protein phosphatase expression (y-axis) vs MoCKa expression (x-axis) in a range of transcriptome datasets from different experiments. (Note: Log2 scale on axes and grids are represented with fixed “aspect ratio” to highlight the different slopes of the correlations). MGG_01690 is not in the pulldown while the other five are and used to illustrate that not all S/T phosphatases are well correlated with CKa. The P values for the Null hypothesis of no correlation with CKa are: MGG_00149 P=2.7E-10, MGG_03154 P=2.5E-8, MGG_06099 P=4.0E-6, MGG_10195 P=6.9E-4, MGG_03838 P=8.8E-4, MGG_01690 P=2.6E-2. Transcriptomic data was downloaded from public websites so as to be able to test the relationship under many different growth conditions in many experiments.

### CKa expression correlates with expression of genes associated with disaggregation and autophagy

Since CK2 activity has the potential to favour protein-protein binding between intrinsically disordered proteins it consequently also has the potential to enhance protein aggregation and aggregation of misfolded proteins. Misfolded proteins or protein aggregates may trigger the unfolded protein response involved in disaggregation. Hsp104 is a disaggregase that cooperates with Yjdg1 and SSa1 to refold and reactivate previously denatured and aggregated proteins (Glover and Lindquist, 1998). Alternatively, protein aggregates that cannot be dispersed by disaggregase may be degraded through autophagy since these kinds of aggregates are too big for proteasome degradation (Wong and Cuervo, 2010). If this is the case, CK2 upregulation should be accompanied by higher autophagy flux or at least there should not be low expression of key autophagy genes when CK2 expression is high (Wong and Cuervo, 2010). ATG8 is a key autophagy protein for which its turnover rate can reflect autophagy flux (Klionsky *et al*., 2016). To test this second sub-hypothesis (Sub-hypothesis 2), we used the expression data we downloaded for plant infection experiments with *M. oryzae* and also for another fungal plant pathogen, *F. graminearum*, that has rich transcriptomic data available (see methods), to examine expression of HSP104, YDJ1, SSA1, and ATG8 relative to CKa. For *M. oryzae*, we found an approximately 60-fold increase in MoHSP104 expression associated with a doubling of MoCka transcript levels. With increasing expression of MoCka the MoHSP104 levels did not increase further. MoSSA1 expression had a similar pattern to MoHSP104 with a 16-fold increase across the initial 2-fold increase in MoCka expression. For MoYDJ1, expression increased with MoCKa expression, but not as dramatically (**Fig. 2**). For *M. oryzae*, we find a log-log linear relationship between the MoCKa expression and MoAtg8 expression (**Fig. 2**). Since CKa expression disaggregase expression and ATG8 expression can be expected to all increase with growth rate these observed relationships could simply be a reflection of increased growth rate if all activities are only growth rate correlated. If that is the case the correlation should become very weak, disappear or become negative if compensated for growth rate. The histones are tightly connected to growth rate since overexpression of histones are cytotoxic (Mei *et al*., 2017). The two rightmost plots (Fig. 2 and Fig. S2) shows that HSP104 and ATG8 well correlated with CKa also when compensated for growth rate indicating that increased disaggregase and ATG8 activity is needed with higher CKa activity and that these relationships are not just growth rate dependent. In the case of *F. graminearum*, we also found increased expression of all of the genes correlated with FgCKa expression across a large range of experiments) (**Fig, S1A, B**). As for *M. oryzae* these correlations were also strong when compensated for growth rate (Fig. S3 and S4). Within the *F. graminearum* experiments a time course experiment was selected to examine expression of these genes during the course of infection. Once again, the relationship could be observed (**Fig. S1C**). Overall, these correlations support the hypothesis that protein disaggregation and autophagy are increasingly needed to remove protein aggregates stimulated to form by increasing levels of CKa and its activity in the cell.

**Fig. 2.**
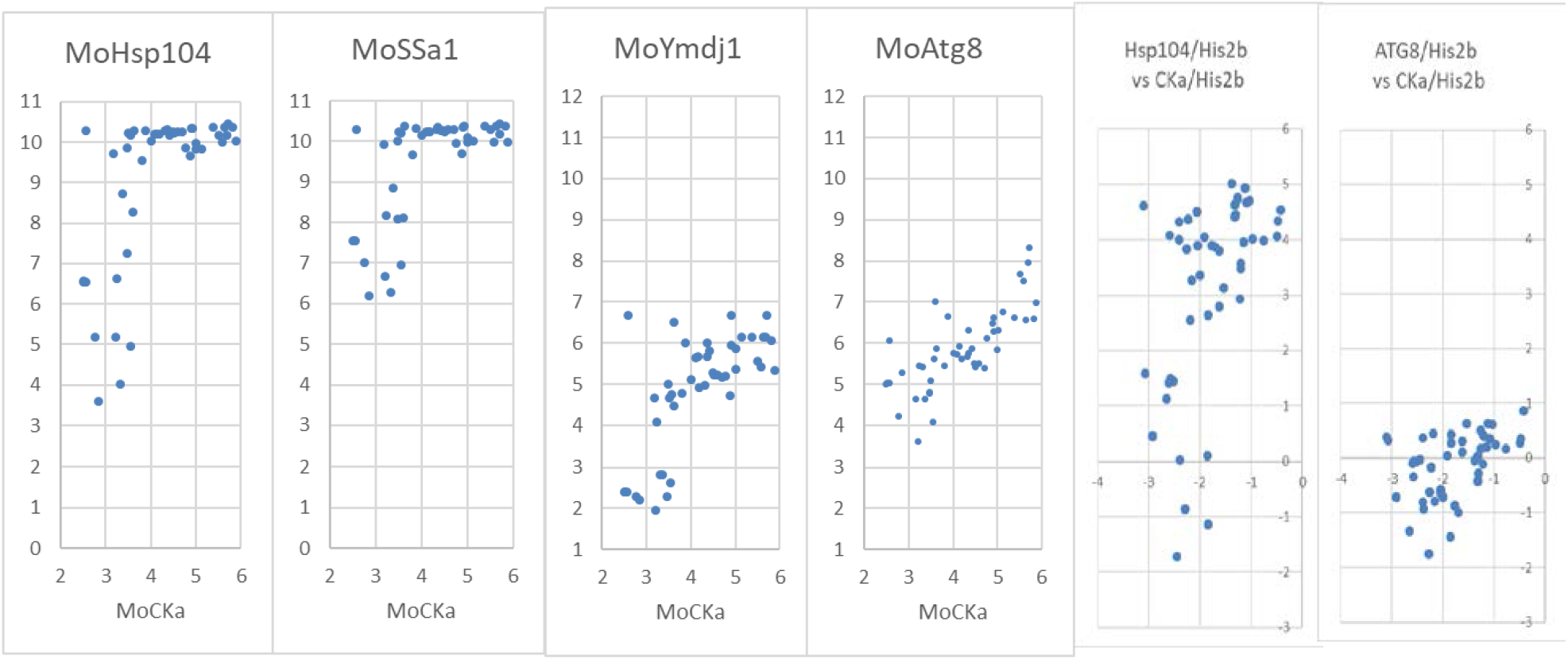
Plot of expression involved in protein quality control vs MoCKa expression in a range of transcriptomes from different experiments (Note: Log2 scale on axes and grids are represented with fixed “aspect ratio” to highlight the slope of the correlation). The two rightmost figures show growth compensated plots using ratio of expression in relation MoHis2b expression that tightly correlates with new DNA synthesis. Hsp104, SSa1 and Ymdj1 has in yeast been shown to cooperatively help aggregate proteins to be able to refold. The key protein with its main function in this process appear to be Hsp104 (Glover and Lindquist, 1998).

In the absence of well-functioning autophagy removing incorrectly formed larger protein aggregates, like those formed in brain cells of Alzheimer’s patients (Zare-shahabadi *et al*., 2015), CK2 activity also facilitates protein aggregate formation and hastens the progression of Alzheimer’s disease (Rosenberger *et al*., 2016) giving further support for a function of CK2 in facilitating the formation of protein aggregates from intrinsically unfolded proteins that are then subjected to autophagy.

Interestingly a specific CK2 phosphorylation motif only present in primates has been shown to be involved in aggressome formation and clearance by autophagy (Watabe and Nakaki, 2011, 2012) pointing to a need for high aggregate clearance activity when CK2 levels are high. As autophagy is important to appressorium development (Liu and Lin, 2008; Kershaw and Talbot, 2009), it will be of interest to further examine the role of the CK2 ring structure during appressorial development and infection (Zhang *et al*., 2019).

However, as infection proceeds in into the necrotrophic phase for *F. graminearum* CKa is progressively upregulated pointing to a role for it in the necrotrophic stage (**Fig. S1C**) but then mainly in probably mainly in defence against plant defences stressing the fungus creating problems with protein folding. The 32.01.07 unfolded protein response was the only sub-category within the main category 32 “CELL RESCUE, DEFENSE AND VIRULENCE” that were overrepresented (p-value for the null hypothesis of same frequency as in the whole proteome 1.81E-11).

As a counterpart to CK2 in gelling/un-gelling, disaggregase activity involving the MoHSP104 complex, may be critical for control. Our observation of transcriptional co-regulation between CKa and HSP104 supports this notion. As MoCK2 is present both in the cytoplasm and nucleoplasm it would generally assist intrinsically disordered proteins in protein interactions (Uversky, 2015). It also seems to be essential for assembling ribosomes containing large numbers of intrinsically disordered proteins (Uversky, 2015). All these functions also explains why CK2 is needed constitutively (Meggio and Pinna, 2003) and we find it relatively highly expressed under all conditions but very high under stressed conditions characterized by high levels of autophagy.

## Conclusion

We conclude that CK2 likely has an important general role in the correct assembly/disassembly of intrinsically disordered proteins in addition to its already suggested role in organelle biogenesis (Rao *et al*., 2011). Our results further point to one of the main functions of the CK2 holoenzyme as a general facilitator of protein-protein, protein-RNA and protein-DNA interactions important for a large range of cellular processes. This includes a potential role for it influencing gel formation that creates membraneless organelles at fungal septa, and maybe other pores between other eukaryotic cells, through its interaction with (and modification of) intrinsically disordered proteins in or around the pore opening. To cite Meggio and Pinna (Meggio and Pinna, 2003) “Using basket terminology, one would say that CK2 looks like a “playmaker” not a “pivot”: hardly ever does it make scores; nevertheless, it is essential to the team game”.

## Data availability

Additional data that support the findings of this study are available from the corresponding authors upon request.

## Acknowledgements

Acknowledgements We thank Dr. Guanghui Wang, Dr. Wenhui Zheng, Dr. Ya Li and Dr. Huawei Zheng (Fujian Agriculture and Forestry University, Fuzhou, China) for their helpful discussions. This work was supported by the National Natural Science Foundation of China (U1805232), National Key Research and Development Program of China (2016YFD0300700) and National Natural Science Foundation for Young Scientists of China (Grant No.31500118 and No.31301630).

## Authors’ contributions

Conceived and designed the study: L. Z., D. Z., Z. W, B. O. H. and S. O. Performed the experiments: L. Z., D. Z., D. L., H. L, Conceived, developed and tested the new methods: S. O. Prepared secondary data for analysis: B. O. H., Analysed the primary data: L. Z., and S. O. Analysed the secondary data: B. O. H. and S.O. Wrote the paper: L. Z., D. Z., Z. W., B. O. H. and S. O.

## MATERIALS AND METHODS

### Material and methods details

#### Pulldown and identification of CKa interacting proteins

*M. oryzae*, strain GFP-MoCKa (GfpA) (Zhang *et al*., 2019), was grown on liquid CM medium consisting of 0.6% yeast extract (CAS No. 8013-01-2), 0.6% casein hydrolysate (CAS No. 65072-00-6), 1% sucrose (CAS No. 57-50-1) in DD-water for 3 days at 25°C in the dark. The cultures were harvested and total protein samples were extracted from vegetative mycelia and incubated with anti-GFP beads (GFP-Trap®_A, Chromotek, Hauppauge, NY, USA) 90 minutes at 4°C with gentle shaking following the manufacturer’s instruction. After a series of washing steps, proteins bound to anti-GFP beads were eluted. The eluted proteins were sent to BGI Tech (Shenzhen, Guangdong province, China) and analysed by mass spectrometry for analysis of sequence hits against the *M. oryzae* proteome. The transformant expressing GFP protein only was used as the negative control and the Ku80 background strain (Zhang *et al*., 2019) was used as Blank control. Data from three biological replicates were analysed against the background of proteins that were bound non-specifically to the anti-GFP beads in GFP transformant and in Ku80 to get the final gene list of genes that was pulldown with MoCKa (Data S1).

### Estimation of non-specific binding of proteins in the pulldown

We developed two methods to estimate the number of non-specific binding proteins found in the CKa pulldown. The first approach is a chemistry-based reasoning and assumes that the degree of unspecific association to the protein per protein surface area is the same for GFP specific hits and for the CK2 holoenzyme pulled down. Using this technique, we estimate that 44-132 proteins are false positive in the CKa pulldown (all proteins pulled down by GFP-Beads or the Beads alone already removed from the list) (**Data S1**). The Second approach is statistical where we assume that binding of the true interacting proteins to CKa are log-normally distributed related to the abundance of each protein in the pulldown, since the median is low and close to zero and negative amounts are impossible. Using the deviation from the theoretical distribution, with higher than expected amounts of a specific protein, for the less abundant proteins we estimate that 46-81 proteins found in the CKa pulldown (with controls subtracted) were false positive. The higher number was used to set a conservative threshold for which proteins should be included in the analysis (See **Data S1** for details of both methods).

### Finding *M. oryzae* proteins containing the helix unfolding motif

The MEME motif LSDDDXE/SLEEEXD (Zetina, 2001) was used to search through the proteome of *M. oryzae* using the FIMO tool at the MEME suite website (http://meme-suite.org/). Results were then downloaded and handled in MS Excel to produce a list of proteins with at least one motif hit (For data see **Table 1**).

### Analysis of CKa expression in relation to disaggregase related protein, Atg8 and Ser/Thr phosphatase expression

For *M. oryzae*, transcriptome experiment data was downloaded as sra/fastq files from Gene Expression Omnibus, https://www.ncbi.nlm.nih.gov/geo/ and mapped onto the genome found at http://fungi.ensembl.org/Magnaporthe_oryzae/Info/Index. The procedure was the following: Gene Expression Omnibus (Barrett *et al*., 2012) was queried for SRA files originating from *M. oryzae* and the files downloaded. The conversion from SRA to Fastq was done using the SRA toolkit (http://ncbi.github.io/sra-tools). The resulting samples were subjected to quality control using FASTQC (Andrews, 2010). Quantification of RNA was performed using Kallistowith default settings (Bray *et al*., 2016), the data was then normalized using the VST algorithm implemented in DESeq2 (Love *et al*., 2014).

*F. graminearum* transcriptomic data (FusariumPLEX) was directly downloaded from the PlexDB database selecting *in planta* experiments and treatments based on the information found in the database (http://www.plexdb.org/modules/PD_general/download.php?species=Fusarium).

For each fungus an expression matrix was prepared with the different experiments as columns and gene IDs as rows, using the MGG codes for M. oryzae and FGSG for *F. graminearum* according to BROAD, respectively. From the resulting matrixes (For these data see Key resource **Table 1**) we used the data needed to plot expression of selected proteins vs CKa expression for the two fungi.

*M. oryzae* data are expressed as log2 of RPKM values. Similarly, for *F. graminearum*, these data were already as log2 of reported relative expression. Gene expression data used were from MoCKa, MoHSP104, FgHSP109, MoYDJ1, FgYDJ1, MoSSA1 and FgSSA1 homologues identified in this study (see results) as well as from, MoCKa (MGG_03696) (Zhang *et al*., 2019), MoAtg8 (MGG_01062) (Veneault-Fourrey *et al*., 2006), FgCKa (FGSG_00677) (Wang *et al*., 2011; Zhang *et al*., 2019), FgAtg8 (FGSG_10740) (Josefsen *et al*., 2012), MoHistone2b (MGG_03578)(NCBI) FgHistone2b (FGSG_11626). Data from the *M. oryzae* expression matrix was also used for plotting MoCKa expression versus the expression of annotated serine/threonine phosphatases found in the CKa pulldown.

## DATA AVAILABILITY

Data and resources external to this paper and data from this study are available as listed in and linked from Table 1.

---

**Fig. S1.**
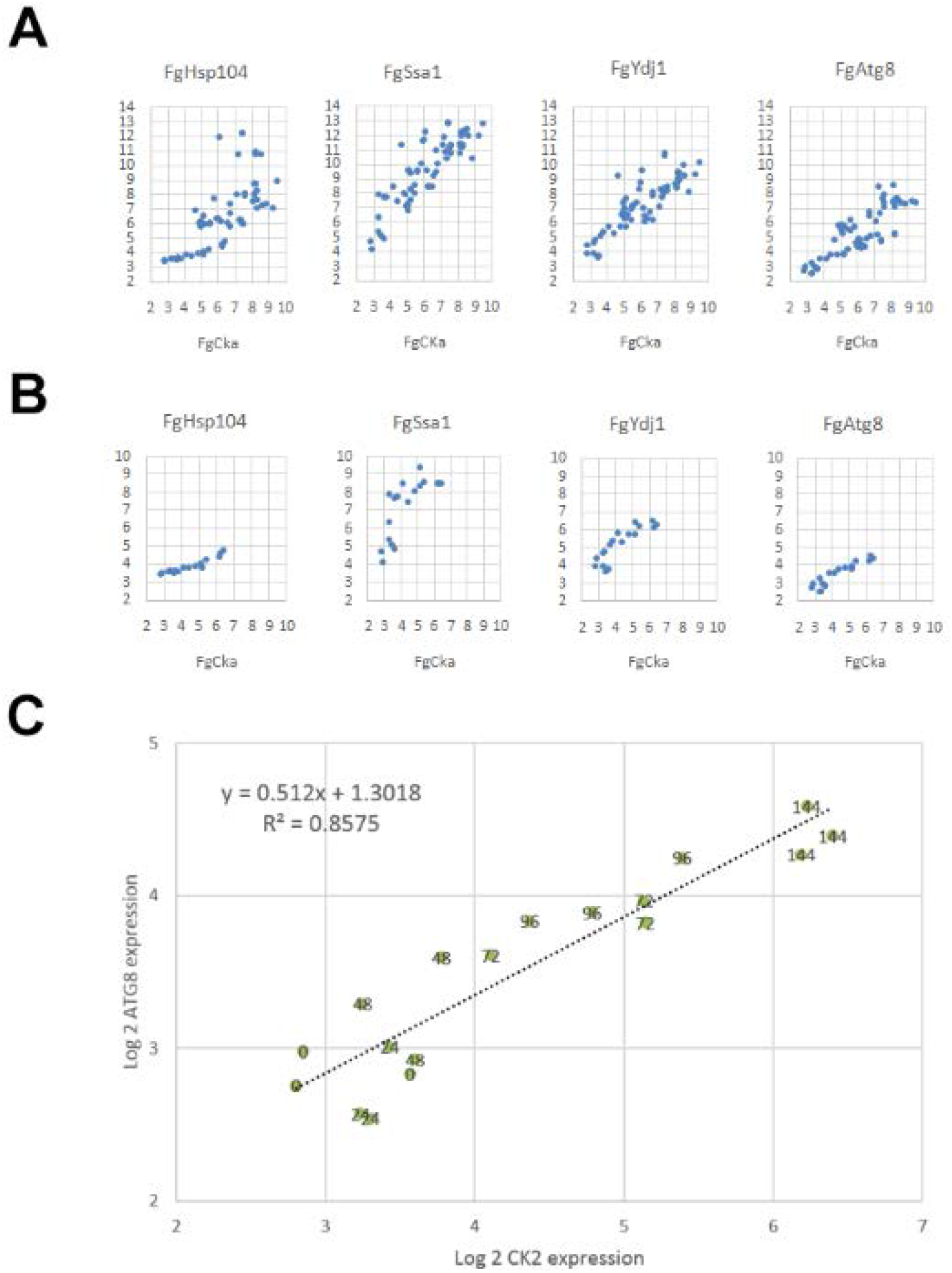
Plot of expression involved in protein quality control vsFgCKa expression in a range of transcriptomes from different experiments. (Note: Log2 scale on axes and grids are represented with fixed “aspect ratio” to highlight the slope of the correlation). **(A)**. data from all experiments. **(B)** data from a time course infection experiment. Hsp104, SSa1 and Ymdj1 have in yeast been shown to cooperatively help aggregate proteins to be able to refold. The key protein with its main function in this process appear to be Hsp104 (Glover and Lindquist, 1998). **(C)** Plot of FgAtg8 (autophagy) expression vs FgCKa expression in a times series infection experiment with 3 replicates where numbers in the plot indicate hours post infection (hpi). (Note: Log2 scale on axes and grids are represented with fixed “aspect ratio” to highlight the slope of the correlation). P value for the Null hypothesis that there is no correlation = 3.6E-08.

**Fig. S2.**
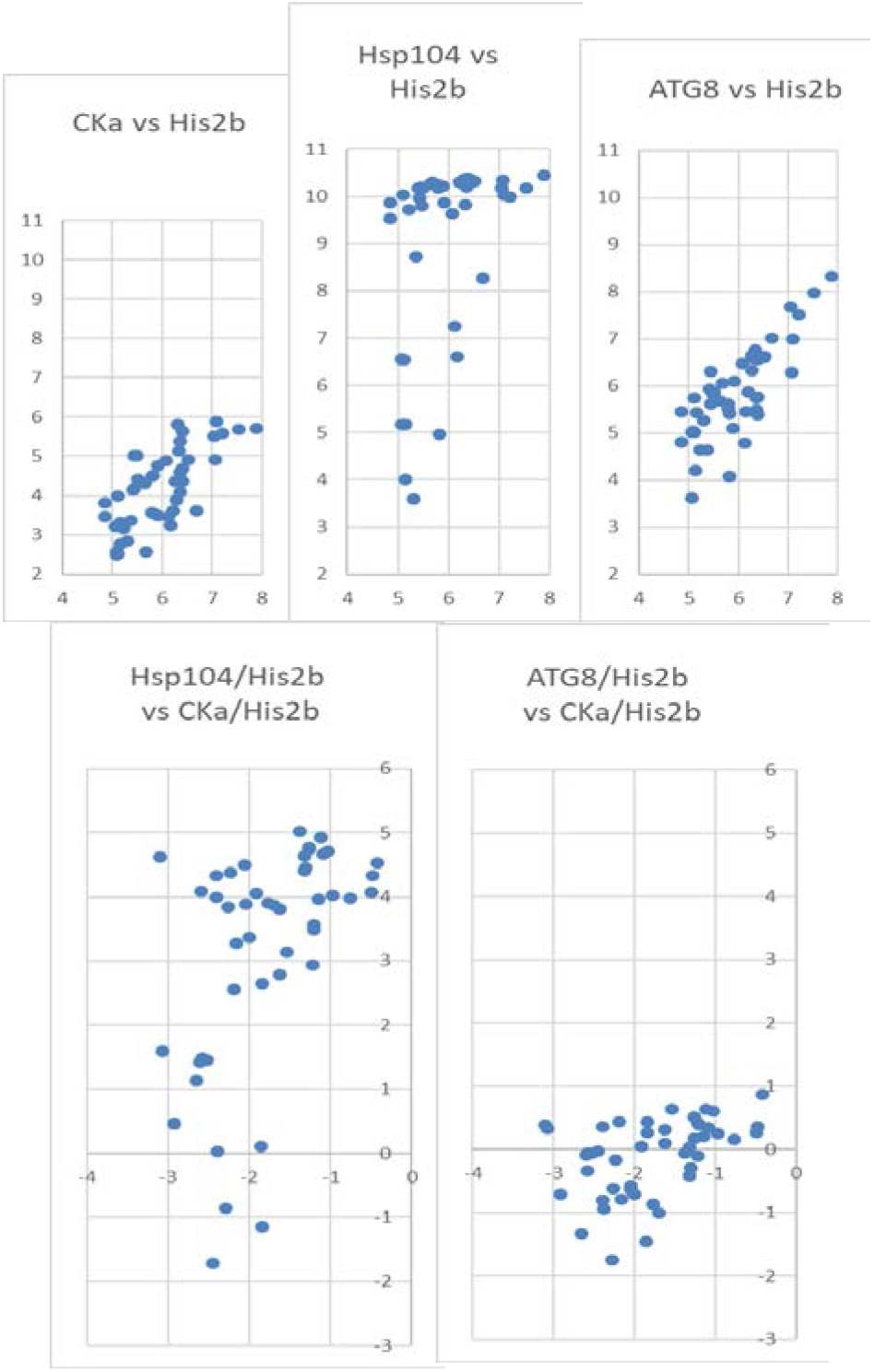
Expression of MoCKa, disaggregase MoHsp104 and MoATG8 in relation to growth as MoHis2b expression. Top 3 diagrams show CKa, the disaggregase Hsp104 and ATG8 expression plotted against growth reflected by His2b expression. The 2 bottom diagrams show growth adjusted plots where the expression of the disaggregase and the ATG8 expression per His2b expression are plotted against CKa/His2b expression as in a qPCR using His2b as reference gene to detect changes in relation to growth. Scales are Log2.

**Fig. S3.**
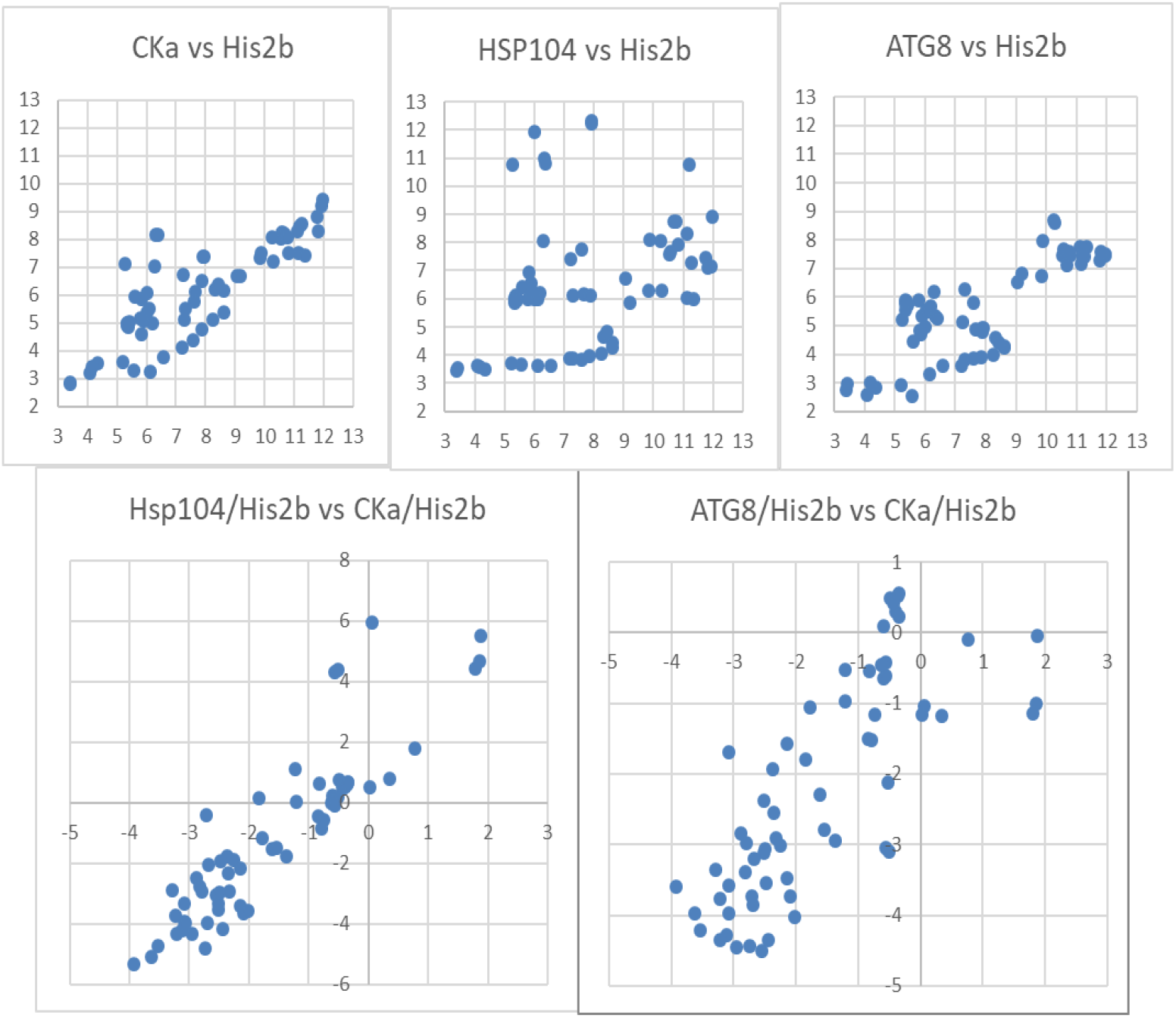
Expression of *Fusarium graminearum* FgCKa, disaggregase FgHsp104 and FgATG8 in relation to growth as FgHis2b expression. Top 3 diagrams show CKa, the disagggregase Hsp104 and ATG8 expression plotted against growth reflected by His2b expression. The 2 bottom diagrams show growth adjusted plots where the expression of the disaggregase and the ATG8 expression per His2b expression are plotted against CKa/His2b expression as in a qPCR using His2b as reference gene to detect changes in relation to growth. Scales are Log2.

**Fig. S4.**
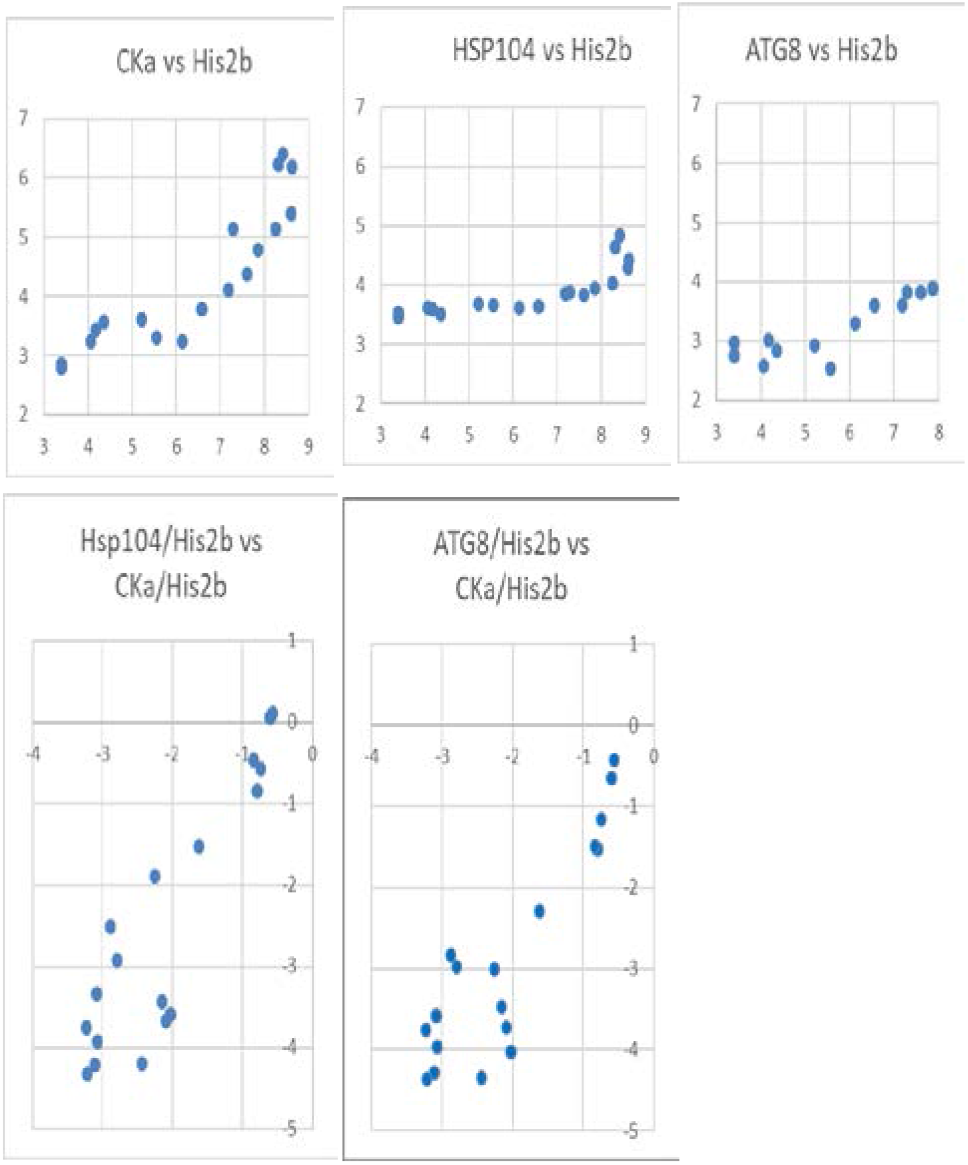
Expression of *Fusarium graminearum* FgCKa, disaggregase FgHsp104 and FgATG8 in relation to growth as FgHis2b expression. Data from one experiment were CKa expression follows hours post infection (HPI) (Fig S1C). Top 3 diagrams shows CKa, the disagggregase Hsp104 and ATG8 expression plotted against growth reflected by His2b expression. The 2 bottom diagrams shows growth adjusted plots where the expression of the disaggregase and the ATG8 expression per His2b expression are plotted against CKa/His2b expression as in a qPCR using His2b as reference gene to detect changes in relation to growth. Top left show that both growth and CKa expression increases with HPI since we previously showed that CKa increases with HPI. The two bottom diagrams shows that the growth compensated relative expression of disaggregase and ATG8 increases with CKa relative expression.

